# Ischaemic Preconditioning attenuates Chronic Renal Damage following Ischaemia Reperfusion Injury

**DOI:** 10.1101/2023.07.13.548869

**Authors:** Charlotte Victoria Maynard Brown, Gilda Pino-Chavez, Aeliya Zaidi, Irina Grigorieva, Emma Woods, Robert Steadman, Rafael Chavez, Soma Meran, Usman Khalid

## Abstract

Acute Kidney Injury (AKI) is a common cause of Chronic Kidney Disease (CKD). The leading cause of AKI worldwide is Ischaemia Reperfusion Injury (IRI), seen most commonly in the clinical setting as a result of sepsis-driven hypotension. We are increasingly recognising, however, that AKI and CKD are one closely associated continuum of disease, rather than distinct entities. Ischaemic Preconditioning (IPC) is a strategy aimed at reducing the deleterious effects of IRI. This study demonstrates an efficacious model of kidney IRI, and the protective influence of IPC in attenuating renal injury/fibrosis.

A rat model of bilateral kidney IRI was used: Male Lewis rats (n=84) were assigned to IRI, sham or IPC. In IRI, renal pedicles were clamped for 45 minutes. IPC groups underwent pulsatile IPC prior to IRI. Kidneys were retrieved at 24-hours, 48-hours, 7-days, 14-days and 28-days, and assessed histologically.

IRI led to marked histological damage and renal fibrosis development by 28 days. Histological injury scores and degree of fibrosis were significantly increased following IRI and attenuated with IPC. IPC resulted in a 66% reduction in renal fibrosis at 28 days (p<0.001). IRI also led to a significant increase in serum creatinine acutely, which was attenuated by IPC (p<0.0001). Interestingly at 14-days, there was limited histological damage and differentiation between IRI and IPC kidneys was difficult.

IPC can protect from both acute and chronic kidney damage. 14-days post IRI represents a transitional phase, which maybe a timepoint for commitment to either fibrosis or recovery, and hence offers potential for therapeutic intervention.

## INTRODUCTION

Renal fibrosis is the histological hallmark of Chronic Kidney Disease (CKD), and Acute Kidney Injury (AKI) is a common cause. AKI is typified by a rapid decline in renal excretory function, with the consequent accumulation of waste products (1). AKI is a clinical term, used to encompass what is pathologically known as Acute Tubular Injury, or ATI. ATI reflects tubular dilatation, loss of brush border, and tubular epithelial cell necrosis (2). Both AKI and CKD are a major problem amongst hospital patients with a huge financial burden for health services (3–5). The leading cause of AKI worldwide is Ischaemia Reperfusion Injury (IRI), seen most commonly in the clinical setting as a result of sepsis-driven hypotension, major surgery or decompensated cardiac failure (1). We are increasingly recognising, however, that AKI and CKD are one closely associated continuum of disease, rather than distinct entities.

Renal fibrosis is the result of dysregulated healing and remodelling that is intended to restore homeostasis following injury. The activation of myofibroblasts, followed by the deposition of extracellular matrix components such as Collagen, Fibronectin, and the glycosaminoglycan Hyaluronan (HA) is a central step in wound healing (6). However, unchecked activation of this process results in excess matrix deposition, disruption of normal tissue architecture and ultimately, fibrosis (7). Identifying ways to reduce ischaemic kidney injury and its resultant AKI and fibrosis, remains paramount in improving patient outcomes. Ischaemic Preconditioning (IPC) is a strategy that prevents the injurious effects of IRI in *in vivo* models. It has, however, proved to be a challenge to translate this into a beneficial therapy in the clinical setting (8). Reasons for this include a limited understanding of its underlying mechanisms, identification of the optimal method of delivering IPC and the heterogeneous nature of the target patient population.

We previously demonstrated the importance of timing and dose-dependence in a protective preconditioning regime that attenuated AKI in a bilateral rat IRI model. We also defined a unique molecular signature that originated from proximal tubular cells (9–11). Here, we use the same IPC regime to characterise its effects on the progression of kidney injury from AKI to the development of renal fibrosis.

## MATERIALS AND METHODS

### In vivo experiments

All animal experiments were conducted according to the United Kingdom Use of Animals (Scientific Procedures) Act 1986, under licences PPL30/3097 and P6B0CD326. Adult (8 to 12 weeks old) male Lewis rats were used (Harlan Laboratories, UK). The rats were housed and handled according to the local institutional policies and procedures licenced by the Home Office. Ethical approval for all the protocols within the Licence was provided by the Animal Welfare and Ethical Review Body under the Establishment Licence held by Cardiff University. The ARRIVE guidelines were followed for all animal experiments.

Analgesia was provided 24h pre-op and up to 1-week post-op in the form of 200mcg of buprenorphine dissolved in 500ml of drinking water. Animals were anaesthetised with Isoflurane. Under aseptic conditions, a midline laparotomy and atraumatic clamping of both renal pedicles for 45 mins was performed followed by reperfusion and recovery for 24h to 28d before terminal anaesthesia. Sham control animals underwent the same operation without the renal pedicle clamping. IPC animals underwent the IPC regime previously described (3 cycles of 2min ischaemia and 5min reperfusion) prior to IRI (9). A schematic of the *in vivo* experiments is shown in Figure 1.

**Figure 1:**
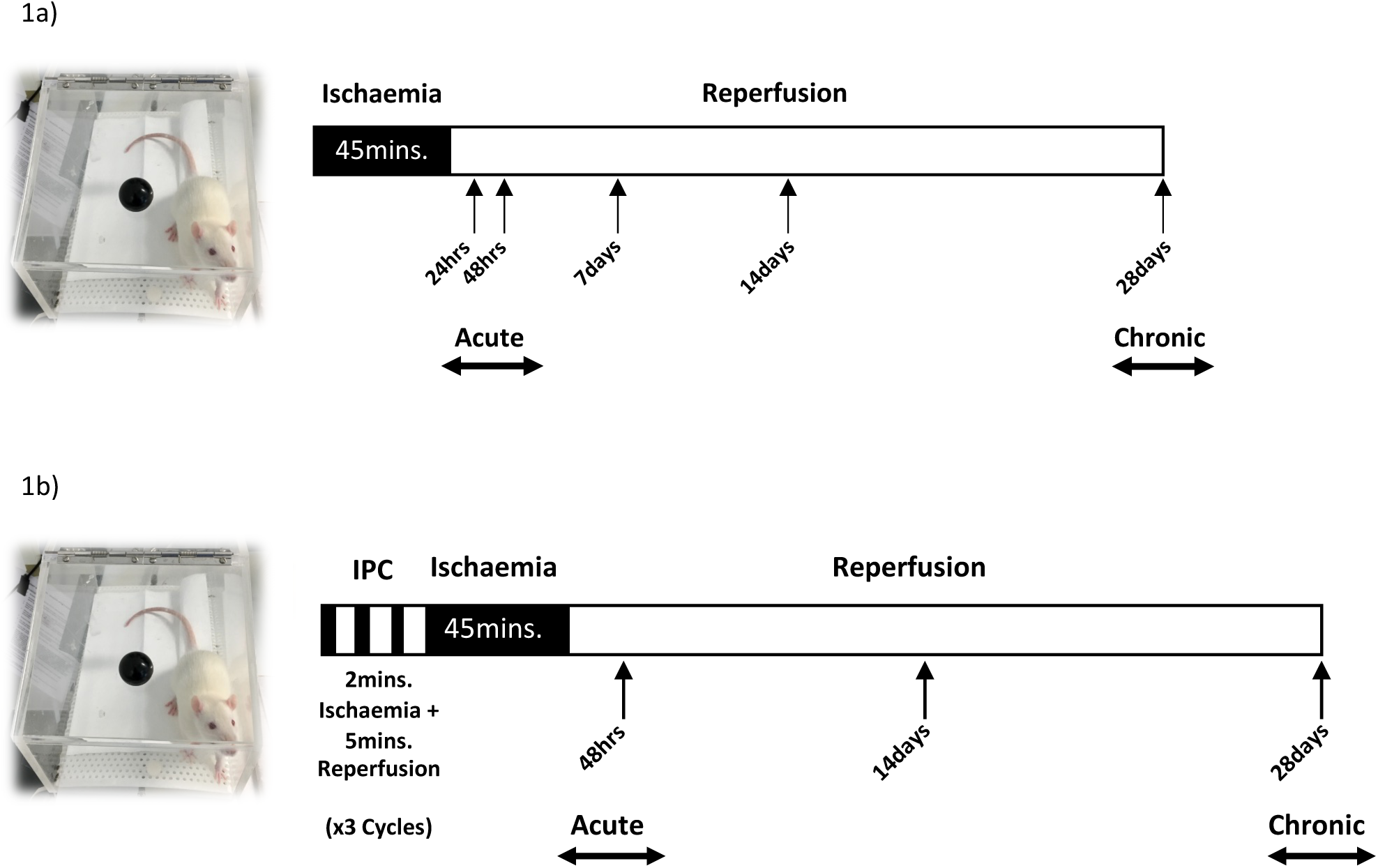
Schematic of animal models a) *IRI model*: adult male Lewis rats underwent a midline laparotomy and were divided into sham operation or bilateral IRI (cross-clamping of both renal pedicles, 45 minutes). These groups were further divided depending on their post-operative observation time: 24 hours (sham n=8, IRI n=8), 48 hours (sham n=8, IRI n=8), 7 days (sham n=4, IRI n=4), 14 days (sham n=4, IRI n=4) and 28 days (sham n=8, IRI n=8). **b) *IPC model:*** IPC+IRI (renal pedicles were identified and cross-clamped bilaterally for 2 minutes (ischaemia), followed by release of the clamps for 5 minutes (reperfusion), performed three times in total, prior to bilateral IRI). IPC+IRI performed only in groups observed for 48 hours (n=8), 14 days (n=4) and 28 days (n=8) post-operatively. In both a) and b): at the designated post-operative time, kidney tissue was retrieved, and blood samples were taken for serum creatinine. Kidneys were retrieved and stored in formalin for histological evaluation.

### Assessment of kidney function

Serum creatinine was measured from blood samples taken at pre-op and at time of retrieval using the Jaffe reaction, according to standard operating procedure at our institution (12).

### Histology

Kidney tissue was embedded in paraffin wax prior to microtomy at a standard thickness of 4μm. Kidney sections were stained with either Haematoxylin and Eosin (H&E) or Masson’s Trichromic according to standard operating procedure. Kidney sections were assessed by an independent experienced histopathologist, who was blinded to the experimental cohort to which the specimen belonged. Acutely injured kidneys (retrieved at either 24 or 48 hours) were assessed using the EGTI scoring system (12), whereas chronically injured kidneys (retrieved at day 7, 14 or 28) were assessed using a novel scoring method (Table 1) by an independent pathologist, who was blinded to the labelling of the tissue specimens. A comprehensive assessment was made regarding the degree of architectural damage observed throughout both the cortex and medulla collectively. Individual scores were assigned for damage in each of the four compartments: tubular, vascular, interstitial, and glomerular. A total score was assigned per kidney section, with a maximum score of 14 and a minimum score of zero.

**Table 1:**
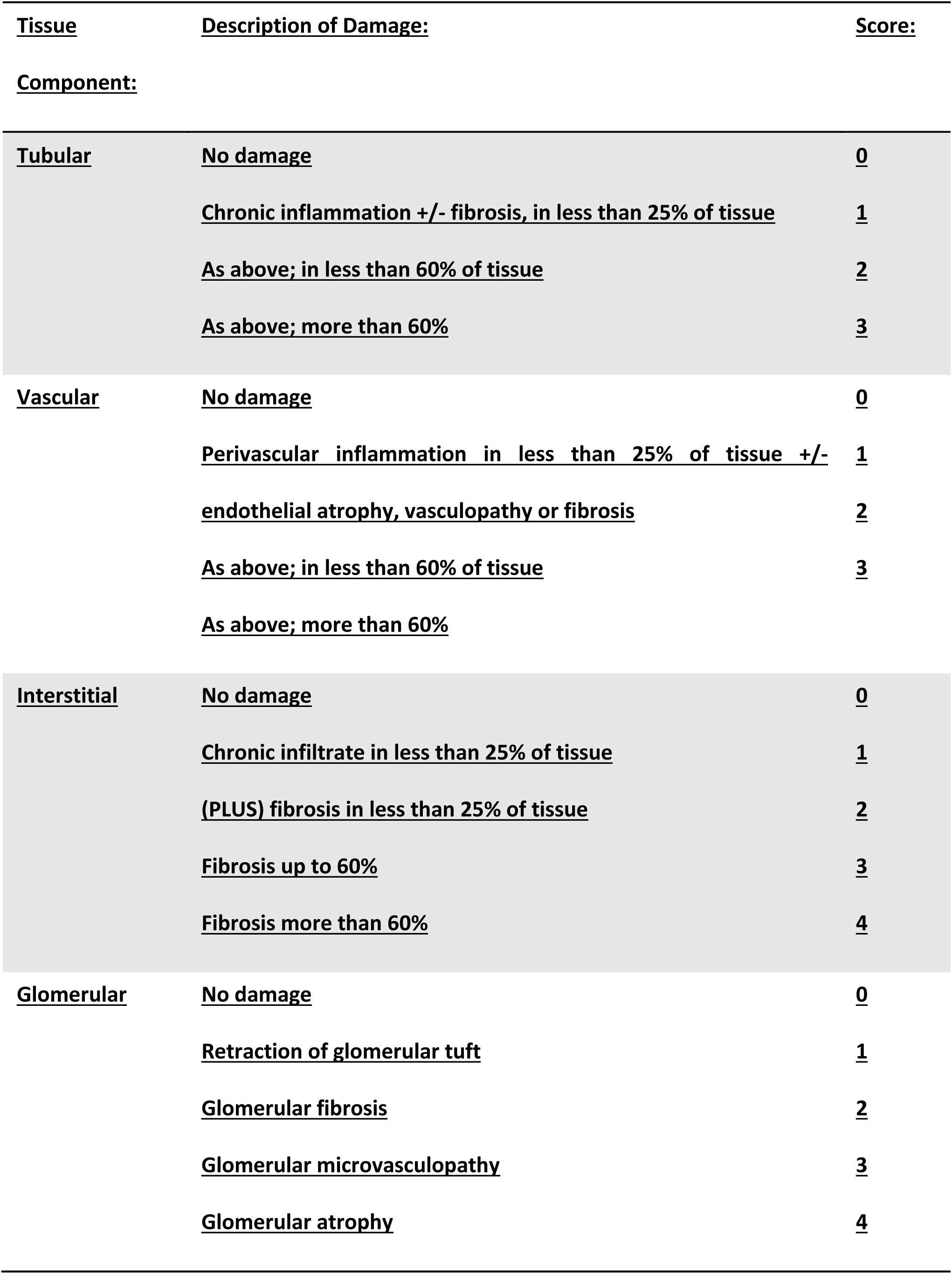
Assessment of histological chronic renal injury. A histology scoring system for chronic renal injury was designed to provide a comprehensive assessment of the degree of architectural damage seen within the kidney, both cortex and medulla, with a cumulative score ranging from 0 to 14. Individual scores were assigned based on the degree of injury in each of the tissue components; tubular, vascular, interstitial and glomerular. Assessment was made by an independent consultant histopathologist, who was blinded to the cohort to which the specimen belonged. Assessment was made using kidney sections stained with Masson’s Trichrome. Left and right kidneys were assessed separately. Median scores were used for statistical analyses.

### Immunohistochemistry

Paraffin-embedded 4μm sections were deparaffinised and autoclaved (Astell AMB240 Autoclave, Liquid Programme) in sodium citrate buffer (10mM, 0.05% Tween, pH 6.0). Blocking of non-specific sites was performed using 10% Normal Goat Serum Blocking Solution (Vector Laboratories).

Primary (Supplementary Table 1) and secondary (supplementary Table 2) antibodies were prepared according to the manufacturer’s recommendation and applied to each section. Vectastain® Elite® ABC Horseradish Peroxidase (HRP) Kit (Vector Laboratories, PK-6100) and DAB Peroxidase (HRP) Substrate Kit (Vector Laboratories, SK-4100) were used prior to dehydration and mounting. All images were obtained using a Leica DMLA Light Microscope. Representative photographs were taken using an Olympus DP27 Microscope Digital Camera.

### Immunofluorescence

Sections were prepared as described above. Primary antibodies (Supplementary Table 3) and corresponding fluorophores (Supplementary Table 4), including 4’,6-Diamidino-2-Phenylindole (DAPI) (ThermoFisher, D1306), were prepared according to the manufacturer’s recommendation and applied to each section. All images were obtained using the Zeiss LSM880 Airyscan Confocal Laser Scanning Microscope.

### Statistical analysis

Statistical analysis was performed using GraphPad Prism Software (Version 8). Data were expressed as either mean ± Standard Error of the Mean (SEM) or median +/-range, depending on whether the data were normally distributed or not. Parametric data were analysed using a t-test (paired or unpaired) or an Analysis of Variance (ANOVA) followed by Tukey’s Post Hoc analysis. Non-parametric data were analysed using a Mann-Whitney U test (unpaired), Wilcoxon test (paired) or Kruskal Wallis test. A p-value <0.05 was considered statistically significant, with levels of significance described in Supplementary Table 5.

## RESULTS

### IRI results in established renal fibrosis

IRI resulted in a significant rise in serum creatinine at 24 and 48 hours post-operatively, with a mean of 162 ± 29.37μmol/L (IRI) compared to 34 ± 0.73μmol/L (sham) (p<0.0001). Serum creatinine normalised in the IRI group by day 7 post-operatively and remained non-significant between IRI and sham groups until day 28 (Figure 2a). Histological injury scores following IRI were significantly higher at 24h (p<0.001), 48h (p<0.0001), 7d (p<0.05) and 28d (p<0.001) post-operatively. There was no significant difference in injury scores at 14 days post-operatively (Figure 2b).

**Figure 2:**
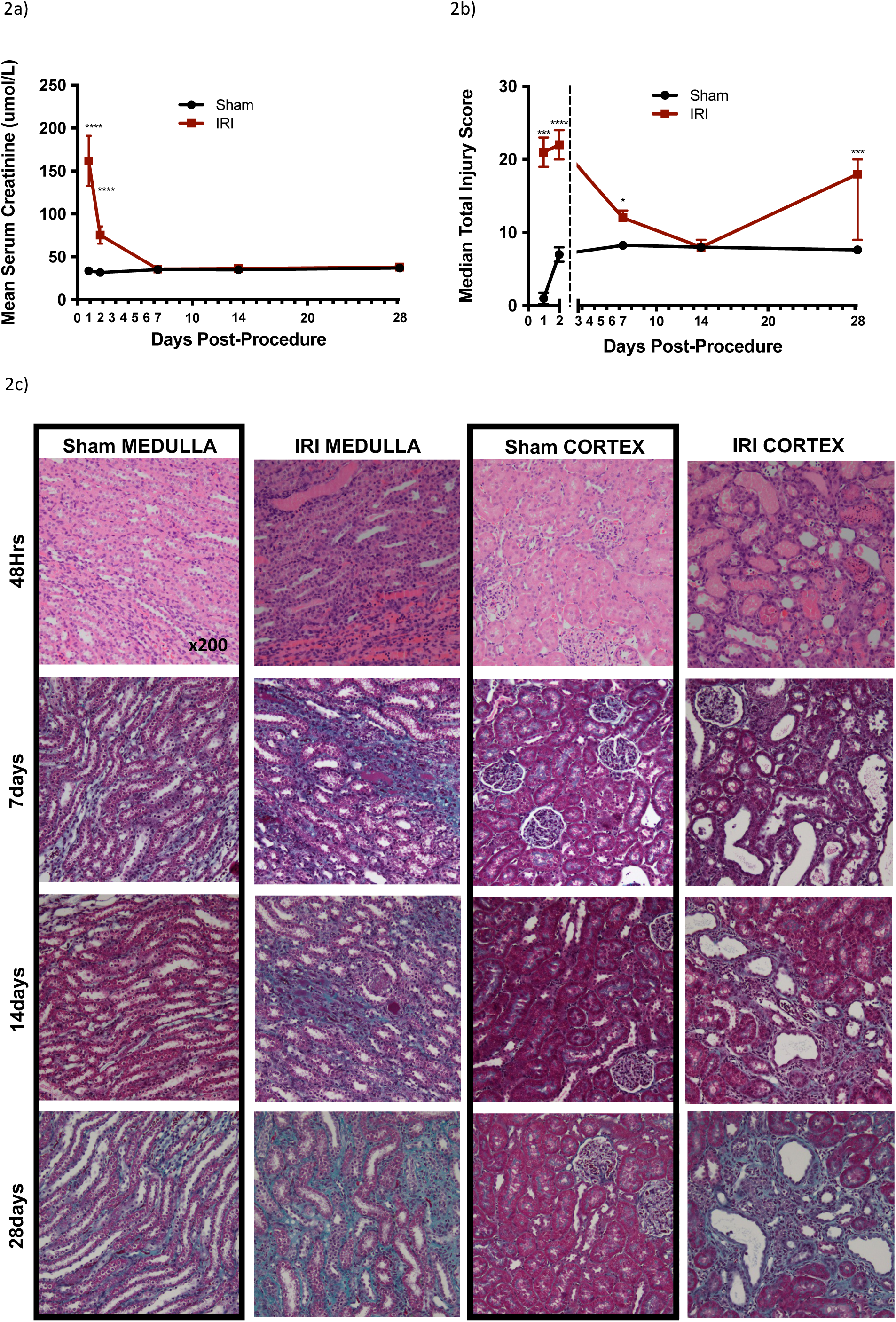
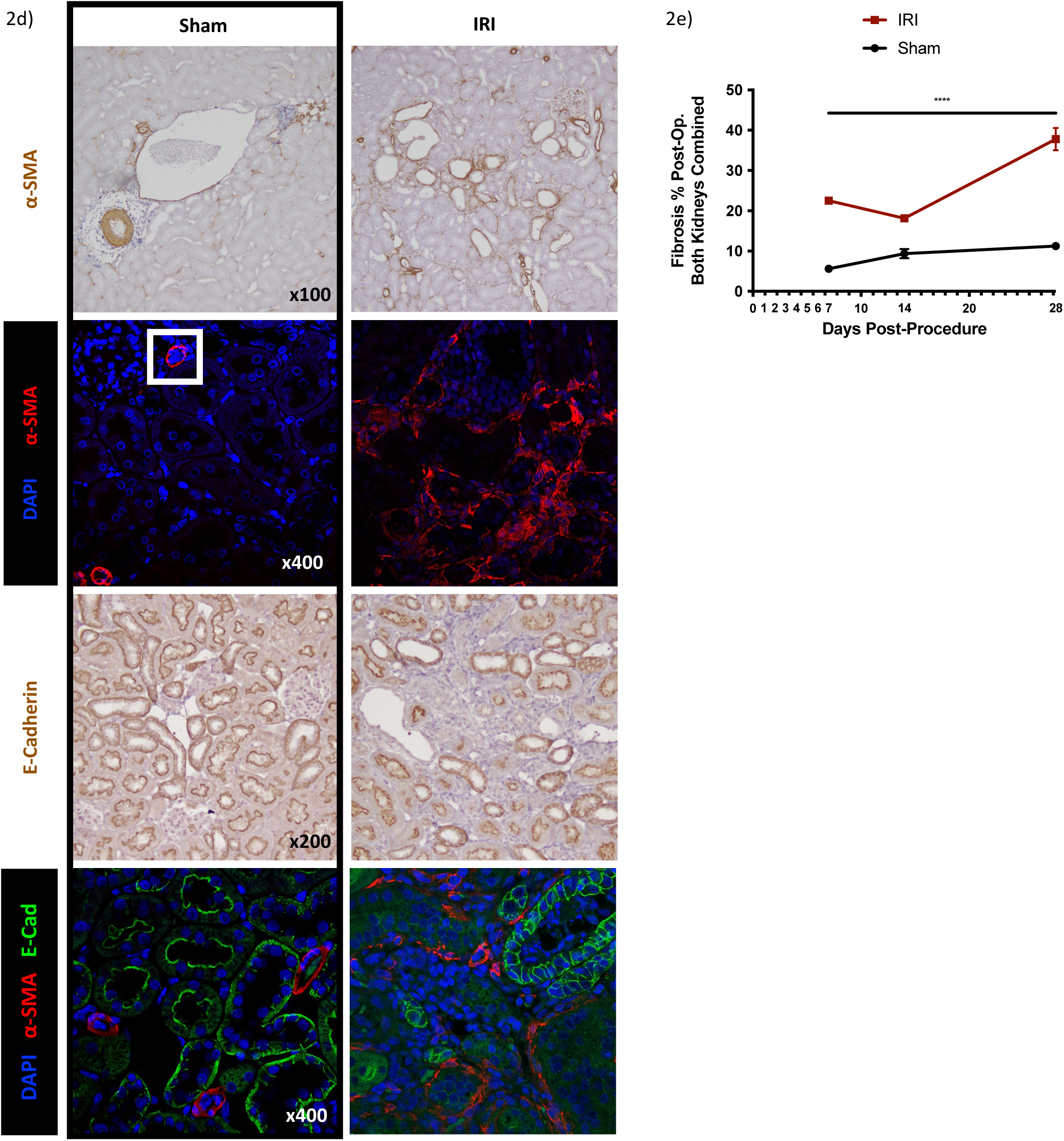
IRI results in established renal fibrosis by 28 days. a) Mean serum creatinine from blood samples taken at times post-operatively (24 hours to 28 days) are shown in both sham (black) and IRI (red) rats. Serum creatinine is measured in µmol and plotted as mean +/-SEM. Statistical significance p**** <0.0001. Statistical significance demonstrated by t-test. b) Median total histological injury scores calculated from whole kidney tissue retrieved at times post-operatively (24 hours to 28 days) are shown in both sham (black) and IRI (red) rats. Histological injury score was calculated using a comprehensive scoring system (Table 2). Median scores were used for statistical analyses and plotted as median + range. Statistical significance p**** <0.0001, p*** <0.001, p* <0.05. Statistical significance demonstrated by Mann-Whitney U test. c) Representative rat renal sections stained with H+E (48 hours) or Masson’s Trichrome (7, 14, 28 days) from sham and IRI rats. d) Representative immunohistochemical and confocal fluorescent microscopic images of rat renal cortex sections retrieved at 28 days post-operatively demonstrating ⍺-SMA (red) and E-cadherin (green). White box in sham cortex demonstrates ⍺-SMA expression in arterial wall. ⍺-SMA expression is used as a marker for myofibroblast presence. Loss of E-cadherin expression is linked to fibrosis expansion. e) Fibrosis percentage observed in both the left, and the right kidney (combined), in IRI rats compared to sham over time post-operatively (7, 14, 28 days).

Histological sections were assessed using Haematoxylin and Eosin (H&E) (48 hours post-operatively), and Masson’s Trichrome (7, 14, 28 days post-operatively). As shown in Figure 2c, IRI resulted in histological damage at all time points. At 48 hours, there was widespread necrosis throughout the cortex and medulla. At 7 days, tubular distortion was observed, with evidence of early fibrosis in the medulla. At 14 days, tubular distortion was less evident than at 7 days, and there was minimal medullary and cortical fibrosis. At 28 days there was widespread fibrosis throughout the cortex and medulla. To confirm the presence of renal fibrosis at 28 days, immunohistochemistry and immunofluorescence were performed using antibodies against α-SMA and E-cadherin (Figure 2d). Interstitial α-SMA was seen following IRI, indicating the presence of myofibroblasts. E-cadherin was observed in distal tubules and collecting ducts in the sham. Following IRI, however, there was a loss of E-cadherin expression in areas of fibrosis. The percentage of fibrosis significantly increased from 23% to 38% at 7 and 28 days respectively (p<0.0001).

### IRI results in differential fibrosis throughout the kidney

To assess fibrosis development, we examined each histological compartment of the kidney individually up to 28 days following IRI (Figure 3). Tubular distortion was marked at 7 days. There was evidence of tubular regeneration at 14 days, with widespread tubular fibrosis by 28 days post-operatively (Figure 3a). Injury to the vasculature was assessed in the same way (Figure 3b). The injury scores were stable, until 28 days post-operatively, when widespread perivascular fibrosis was evident. The interstitium (Figure 3c), showed a widespread inflammatory infiltrate at 7 days, with partial resolution of this at 14 days. By 28 days post-operatively, however, there was marked interstitial fibrosis. Glomerular injury was less evident until 28 days post-operatively (Figure 3d), when glomerular fibrosis and atrophy were apparent. Overall, it was evident that there was markedly less fibrosis at 14 days both within the tubular interstitium, the perivascular areas and the glomeruli with evidence of attenuated inflammation and structural abnormality.

**Figure 3:**
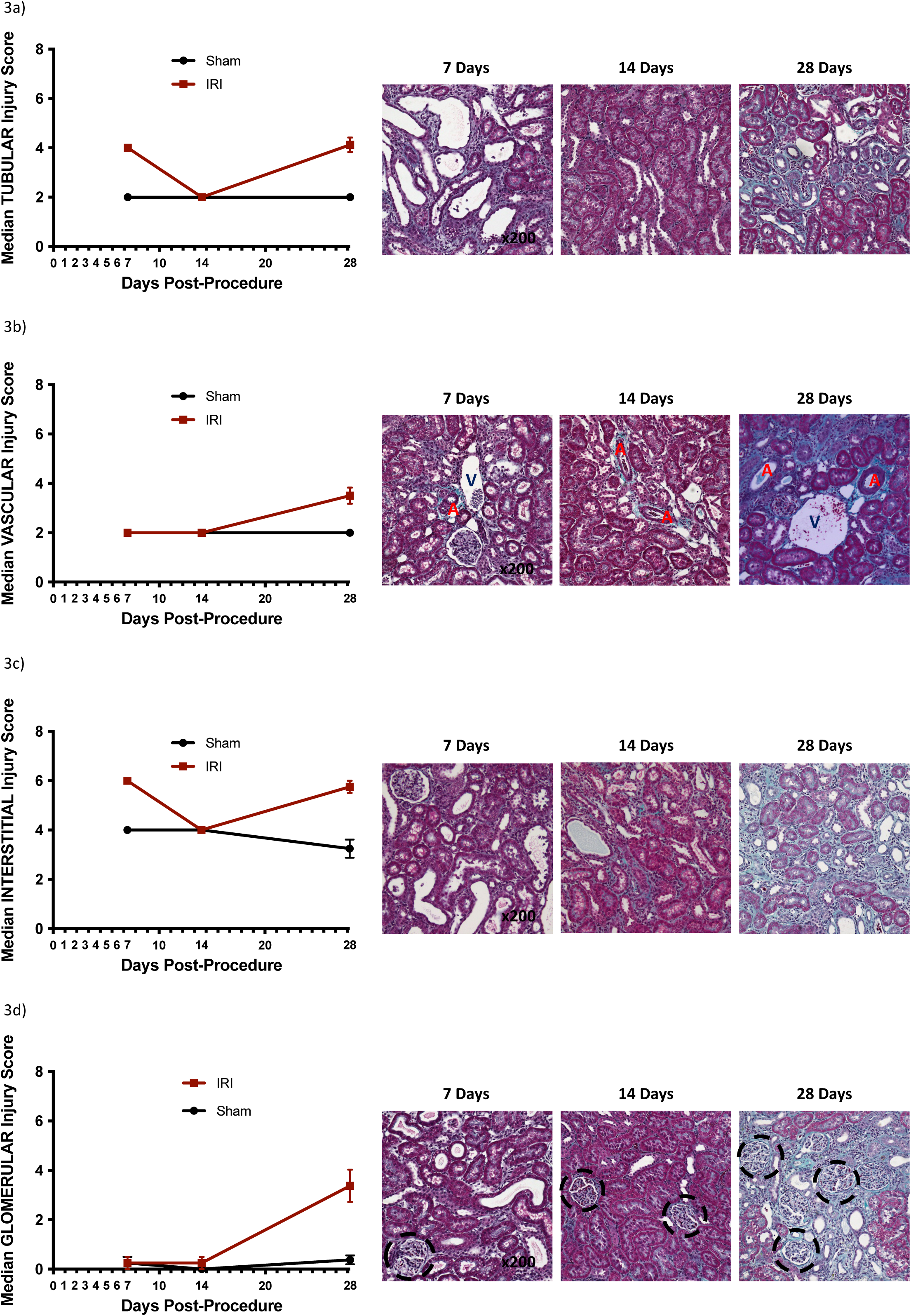
Fibrosis develops across all tissue compartments following IRI. Median histological injury score (sham = black, IRI = red) and representative rat renal sections stained with Masson’s Trichrome are shown in the TUBULAR (a), VASCULAR (b), INTERSTITIAL (c) and GLOMERULAR (d) compartments at times post-operatively (7 [sham n=4, IRI n=4], 14 [sham n=4, IRI n=4] and 28 days [sham n=8, IRI n=8]). [A denotes arterial, V denotes venous structure]. Histological injury score was calculated using a comprehensive scoring system (Table 2). Assessment was made by an independent consultant histopathologist, who was blinded to the cohort to which the specimen belonged. Left and right kidneys were assessed separately, and then total scores illustrated.

### IPC results in protection against fibrosis development

To investigate the protective effect of renal IPC against the development of fibrosis, we applied IPC to our established model of renal fibrosis and performed serum creatinine and histological evaluation (Figure 4). IPC prevented the significant (P<0.001) rise in serum creatinine observed at 48 hours following IRI (Figure 4a), with a mean creatinine of 40 ± 1.59μmol/L in IPC compared to 75 ± 10.18μmol/L in IRI. IPC significantly (P<0.01) attenuated the histological injury measured at 48 hours and 28 days post-operatively, when compared to IRI alone (Figure 4b). Notably, there was no significant difference between the injury scores at 14 days post-operatively in sham, IRI or IPC cohort. H&E and Masson’s Trichrome stains were used to assess histological changes (Figure 4c). IPC attenuated the degree of inflammatory infiltrate observed at 48 hours post-operatively. At 14 days the amount of fibrosis was less in the IPC cohort when compared to IRI alone and this finding was more pronounced at 28 days post-operatively.

**Figure 4:**
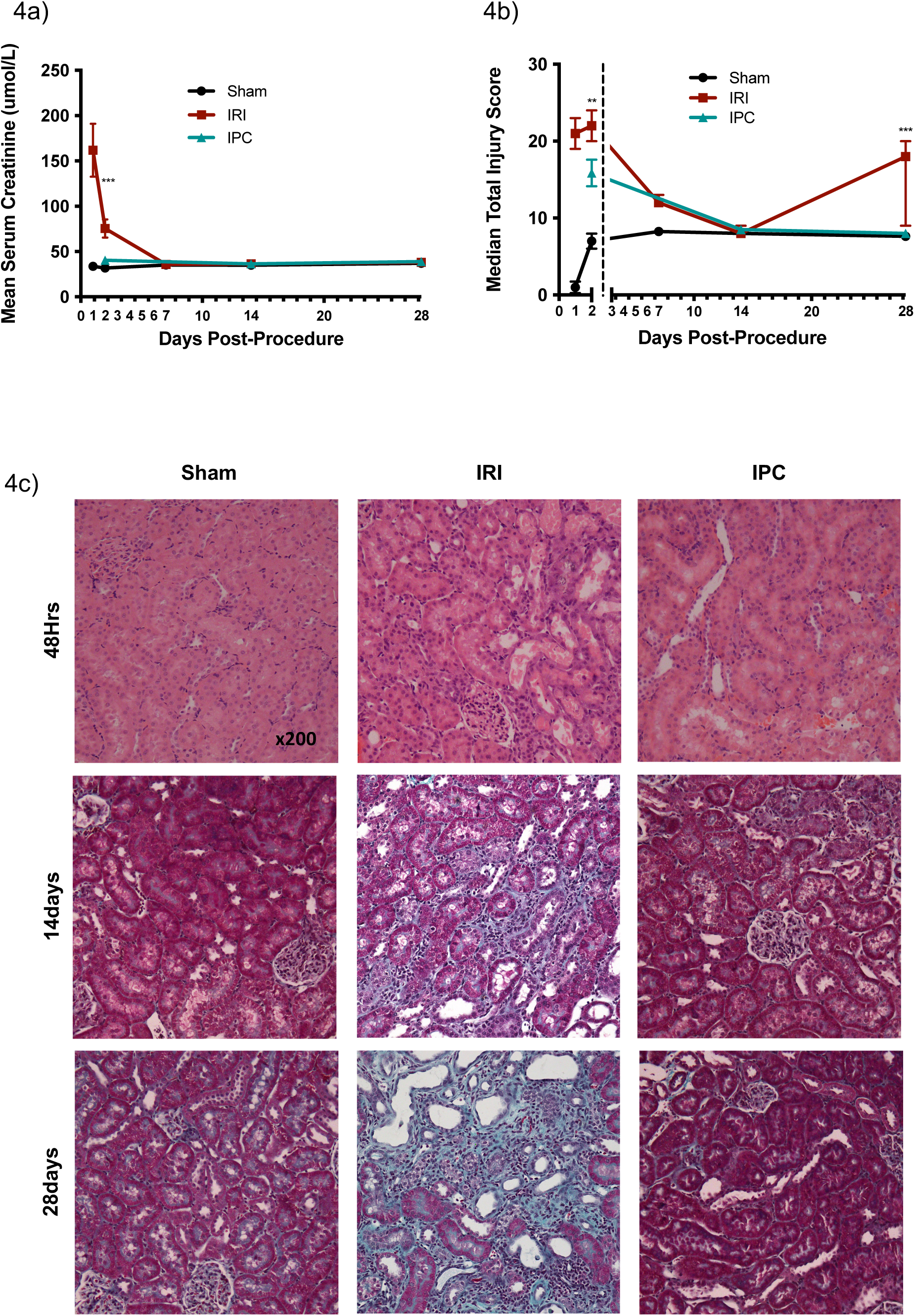
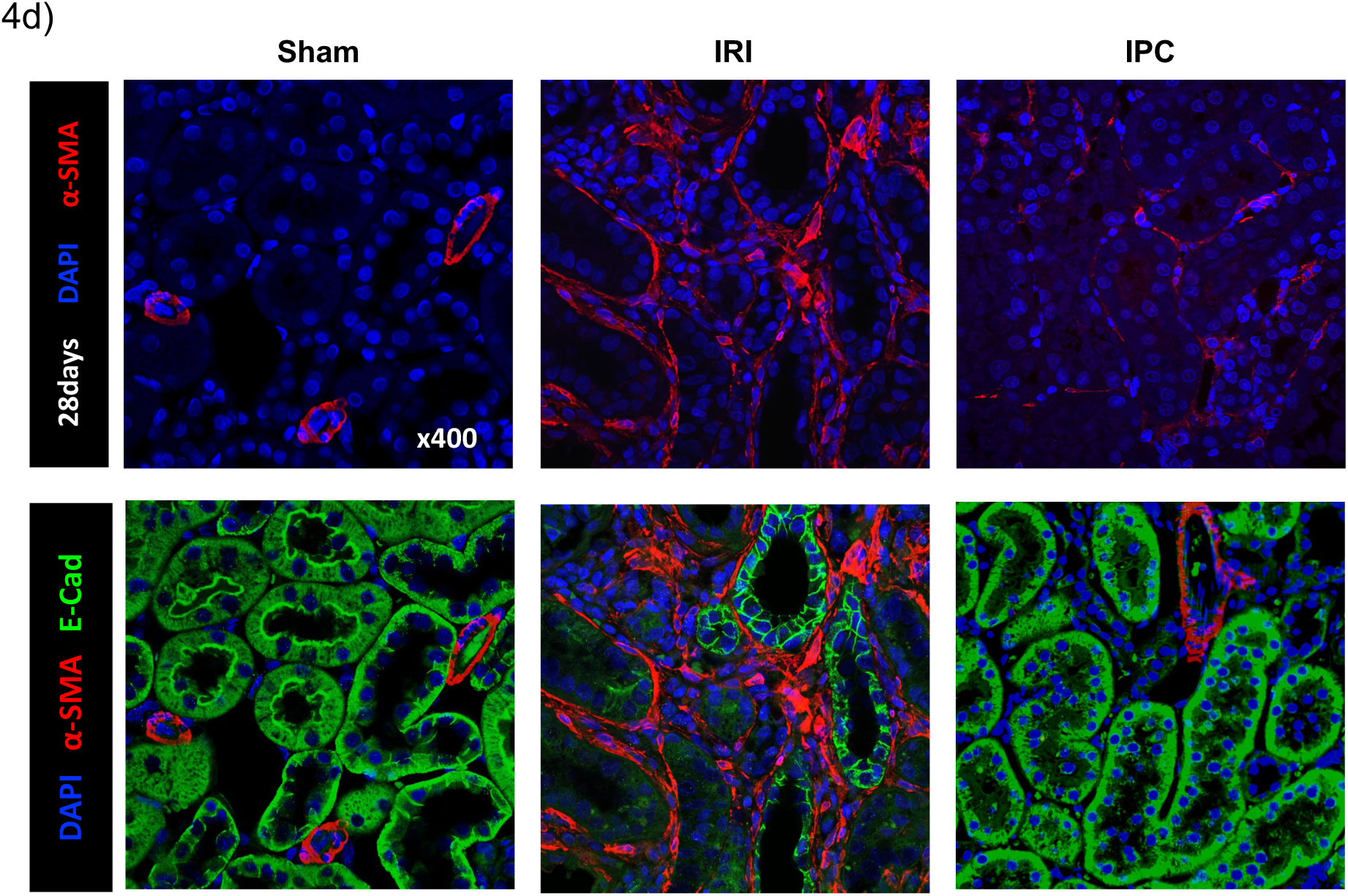
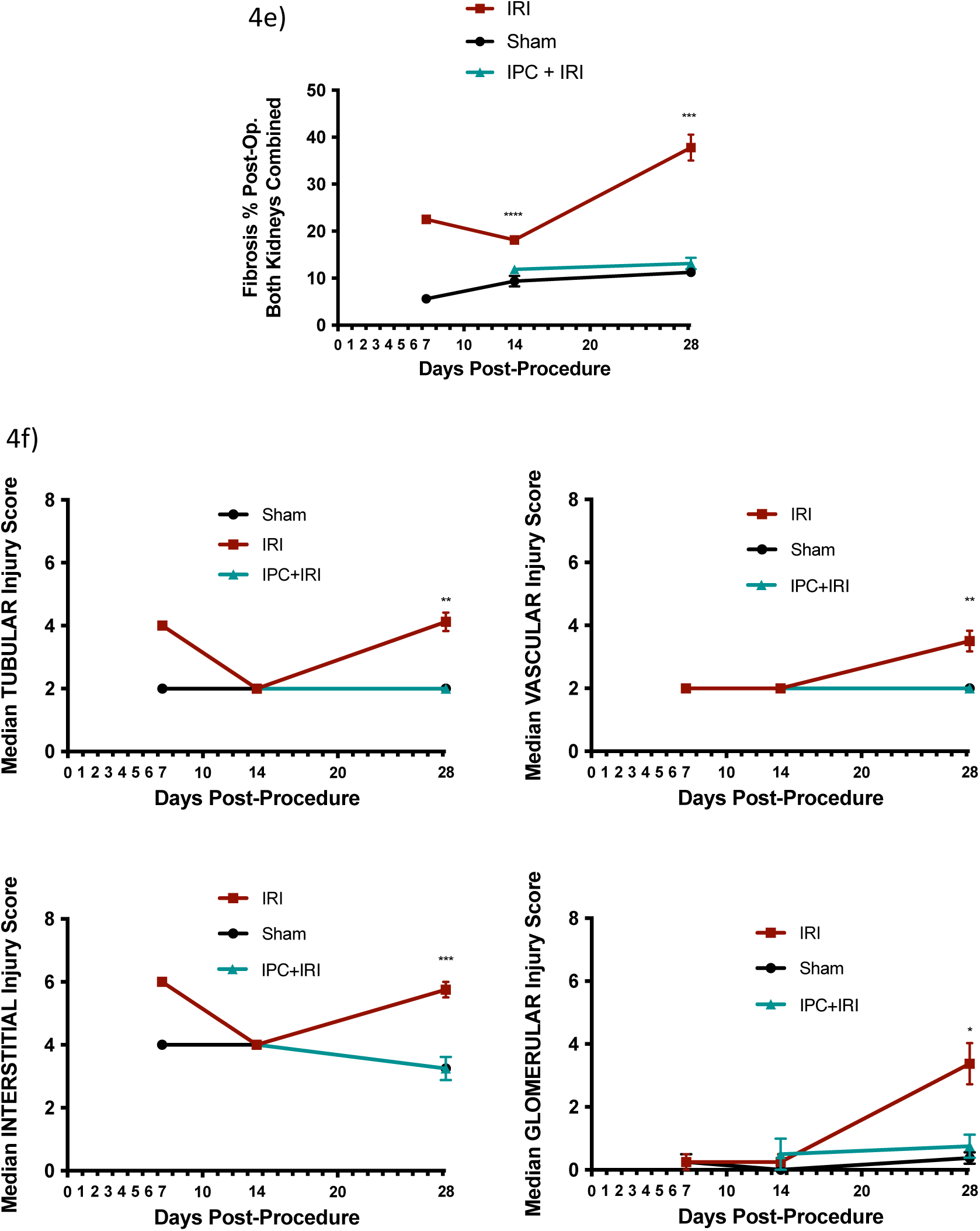
Ischaemic preconditioning inhibits renal fibrosis development. a) Mean serum creatinine from blood samples taken at varying times post-operatively (24 hours to 28 days) are shown in sham (black), IRI (red) and IPC (teal) rats. Serum creatinine is measured in µmol and plotted as mean +/-SEM. Statistical significance (IRI+IPC) p*** <0.001. Statistical significance demonstrated by t-test. b) Median total histological injury scores calculated from whole kidney tissue retrieved at varying times post-operatively (24 hours to 28 days) are shown in both sham (black), IRI (red) and IPC (teal) rats. Histological injury score was calculated using a comprehensive scoring system (Table 2). Assessment was made by an independent consultant histopathologist, who was blinded to the cohort to which the specimen belonged. Left and right kidneys were assessed separately. Median scores were used for statistical analyses and plotted as median + range. Statistical significance (IRI+IPC) p*** <0.001, p** <0.01. Statistical significance demonstrated by Mann-Whitney U test. c) Representative rat renal cortex sections stained with H+E (48 hours) or Masson’s Trichrome (14, 28 days), and representative confocal fluorescent microscopic images of rat renal cortex demonstrating ⍺-SMA expression, from sham, IRI and IPC rats. ⍺-SMA expression is used as a marker for myofibroblast presence. d) Fibrosis percentage observed in both the left, and the right kidney (combined), in sham, IRI and IPC+IRI rats over time post-operatively (7, 14, 28 days). e) a) Median histological injury score in the TUBULAR, VASCULAR, INTERSTITIAL and GLOMERULAR compartments at times post-operatively (7 [sham n=4, IRI n=4], 14 [sham n=4, IRI n=4, IPC+IRI = 4] and 28 days [sham n=8, IRI n=8, IPC+IRI = 8]) are shown in both sham (black), IRI (red) and IPC+IRI (teal) rats.

To confirm the findings observed in histological analysis, immunohistochemistry, and immunofluorescence assessment of α-SMA and E-cadherin expression were performed (Figure 4d). The presence of interstitial α-SMA in the IPC cohort was reduced, when compared to IRI, with only minimal α-SMA evident in the interstitium. In addition, the loss of E-cadherin expression in fibrotic regions of the IRI kidneys was not seen in the IPC cohort. The percentage fibrosis was significantly reduced at both 14 days and 28 days post-operatively in the IPC cohort, from 18% to 12%, and from 38% to 13% respectively (p<0.001) (Figure 4e).

To assess for differential protection offered by IPC against the development of fibrosis, we examined each component of the kidney individually at 28 days following IPC (Figure 4f). Injury scores following IPC mirrored those of sham controls in each tissue compartment, with notably reduced injury scores in the tubular and interstitial compartments at 7 and 28 days compared to IRI alone. Vascular and glomerular injury scores were only seen to increase in IRI at 28 days post-operatively, and this effect was attenuated by IPC.

## DISCUSSION

IPC is a well-documented means of ameliorating IRI *in vivo*, but underlying mechanisms are not well understood and its successful translation to clinical practice remains unresolved. Here we present an established model of IPC and document its effects on the fibrotic process in the kidney, including differential development across tissue compartments over time.

The IRI rat model presented here is a well-established in-vivo model of kidney injury, producing a significant, but not fatal ischaemic injury after 45 minutes of bilateral cross-clamping of the renal pedicles (10, 12–14). The ischaemic insult led to clear histological changes, which resulted in the establishment of renal fibrosis by day 28 post-operatively. This agrees with previous studies (15, 16), demonstrating a robust model of progressive chronic renal histological changes following IRI.

The significant increase in serum creatinine observed at 24 hours post-IRI is consistent with the literature surrounding similar murine models of IRI, as is the rapid normalisation despite the development of chronic histological change (17, 18). Serum creatinine is seen to increase more than threefold (p<0.001) by 24 hours post-IRI, however, this has already started to decrease by 48 hours post injury. Between 48 hours and 7 days post-IRI, the creatinine normalises. The normalisation in serum creatinine over the first 7 days post-IRI demonstrates that development of chronic renal injury cannot be defined on functional assessment alone.

Multiple histological scoring systems exist for the many clinical conditions that result in chronic renal insufficiency, including the Banff working classification of kidney transplant pathology (19) and the Oxford classification of IgA nephropathy (20). However, no existing system could be used to adequately describe the changes associated with this IRI model of evolving chronic impairment. A novel method of characterising the development of chronic renal disease histologically was developed, to include not only global fibrosis, but fibrosis across all tissue compartments. Whilst widely acknowledged to be the key structural component of chronic kidney disease, ambiguity remains about how best to measure renal fibrosis. The degree of fibrosis as a distinct entity is not itself the most useful marker of disease. We are now appreciating more the importance of the distribution of fibrotic changes and their implications. The scoring system presented accounts for fibrosis development in individual tissue compartments: tubular, vascular, interstitial, and glomerular. Not only does this allow more accurate assessment of overall fibrosis, but additionally this enables further understanding of the implications of fibrosis in each compartment. The significant increase in perivascular fibrosis observed when histological damage is greatest at day 28 post-IRI is of interest. Pericytes are a heterogeneous population of cells found in the walls of blood vessels and are thought to contribute significantly to the myofibroblast pool following injury. Lin et al. hypothesised that vascular injury was the most likely precipitating factor in pericyte migration and differentiation to myofibroblast phenotype (21). This could explain the importance of perivascular fibrosis in this model of ischaemic injury, where significant perivascular fibrosis is seen following ischaemic injury once commitment to a fibrotic phenotype is established. Finally, the scoring system accounts for medullary fibrosis, rather than cortical disease alone. This led to the appreciation of earlier onset fibrosis in the medulla over the time course of developing chronic kidney damage and could be important in furthering our understanding of the drive to CKD development.

Injury scores across all four compartments were significantly higher in the IRI group compared to sham. The appearance of α-SMA at 28 days further supported the model of fibrosis development. E-cadherin also represented a marker of renal tubular damage in this IRI model of injury. E-cadherin is a cell adhesion molecule, known for its role in the development and maintenance of renal epithelial polarity (22). Appropriate epithelial polarity is required for normal cell function. In adult kidney, E-cadherin is one of the most abundant cadherins, primarily and reliably expressed in the distal tubules and collecting ducts. Ischaemia is a significant factor which can affect renal epithelial polarity (23), and the changes observed in this model of injury support this, with the observation of loss of E-cadherin expression in those most severely damaged areas following IRI. Epithelial-Mesenchymal Transition (EMT) is considered an important mechanism for the appearance of myofibroblasts and subsequent fibrosis following injury (24). Epithelial cells transitioning to fibroblast phenotype lose E-cadherin expression and consequently cell polarity (25). E-cadherin expression is considered an archetypal cell marker of EMT (25) and its loss of expression in regions of fibrosis development are strongly supportive of this as an appropriate model to study evolving chronic kidney damage.

This 28-day model of IRI shows evolving changes across both the renal medulla and cortex as fibrosis develops. Early changes include distortion of tissue architecture, with tubular necrosis being the key feature acutely, associated with raised serum creatinine. One of the first areas where fibrosis is seen to develop is the perivascular region of the medulla, before involvement of the peritubular interstitium in both medulla and later cortex. Perivascular fibrosis is the deposition of connective tissue components around blood vessels (26). Pericytes are a heterogeneous population of cells found in the walls of blood vessels. In the kidney, pericytes account for nearly a third of all cells in the total tissue and are involved with glomerular filtration as mesangial cells (27). Pericytes are thought to contribute significantly to the myofibroblast pool following injury, and it is thus logical that fibrosis should first initiate in the perivascular location followed by the peritubular interstitium where renal pericytes reside around the peritubular capillaries of the nephron. Lin et al. hypothesised that vascular injury was the most likely precipitating factor in pericyte migration and differentiation to a myofibroblast phenotype. This concept could explain the importance of perivascular fibrosis in this model of ischaemic injury, which can be extrapolated to IRI and AKI in general (21). Unfortunately, this is not reflected in the compartmental injury scores presented here, and the limitations of this are discussed below.

### IPC prevents the formation of renal fibrosis following IRI

Pulsatile IPC prior to IRI resulted in renal protection; not only in preventing AKI (as previously demonstrated), but in preventing fibrosis development and CKD. IPC was evaluated at 48 hours, 14 days, and 28 days post-operatively. Total injury scores at 48 hours and 28 days post-operatively were significantly reduced in the IPC group, when compared to the IRI group alone. There was no difference between the three cohorts at 14 days. At 28 days, injury scores in each of the four tissue compartments were significantly reduced in the IPC group, when compared to the IRI group alone. Histologically, there was less inflammatory infiltrate and tubular distortion in the IPC group at 48 hours post-operatively when compared to the IRI group alone, and this resulted in less fibrosis at 28 days. This protective effect of IPC was observed throughout the follow-up process, suggesting that preventing AKI is a critical factor in averting the development of renal fibrosis.

### Fourteen days post-operatively may represent an integral time-point in our understanding of evolution of fibrosis development versus recovery

A potential point at which the fate of the kidney maybe decided is at the 14-day time point post-operatively. Tubular distortion was evident at 14 days, although not to the degree observed at 7 days post-IRI. Despite there being slightly higher levels of fibrosis evident at this timepoint post-IRI, particularly in the medulla, the injury scores between the cohorts are indistinguishable from one another. This may represent a period of transformation, whereby renal tubules are either regenerating, or committing to irreversible fibrosis. It can be hypothesized that initial tubular necrosis results in significant tubular distortion by day 7 post-IRI. By day 14 this has evolved into either regeneration of those tubules with reparative potential, or the commitment to the development of interstitial and tubular fibrosis. By day 28 there is an irreversible deposition of fibrosis, which could potentially result in renal dysfunction and a rising serum creatinine if the model was left to perpetuate. The 14-day timepoint may represent an important time for pharmaceutical therapeutic intervention; where fibrosis has not yet developed, but commitment to fibrosis may occur. This could offer a timely opportunity to intervene to prevent progression of the IRI group to fibrosis.

In conclusion, we have characterised injury in a rat model of IRI from AKI at 24h-48h to established renal fibrosis by 28 days. Moreover, we have demonstrated that IPC delivered in a pulsatile fashion attenuates this injury and development of fibrosis. Interestingly at the mid-way point (day 14), histological injury scores are similar in preconditioned and injured kidneys, and this maybe a critical point at which the kidneys are initiating the process of recovery or development of fibrosis. Further transcriptomic analyses will help delineate the underlying molecular pathways which maybe contributing to this and potentially open up an exciting avenue for potential intervention.

## ACKNOWLEDGEMENTS

The authors would like to acknowledge the Wales Kidney Research Unit (WKRU), Medical Research Council (MRC), Health and Care Research Wales, and Cardiff University.

## AUTHOR CONTRIBUTIONS

CVMB carried out the animal experiments, the IHC experiments, analyses and wrote the first draft of the manuscript. GPC designed the histology scoring system and carried out the scoring of H&E sections in a blinded fashion. AZ carried out some of the IHC experiments. IG and EW oversaw the IHC experiments and analysis. RS and RC provided intellectual input. SM and UK directed the study conception and design, provided the intellectual input, were involved in data analysis and revised the manuscript. All authors approved the final manuscript.

